# A 3D Brain Geometry Toolkit for Multisite Neuroimaging Analysis

**DOI:** 10.64898/2026.06.29.733626

**Authors:** Yanghee Im, Melody J.Y. Kang, Boris A. Gutman, Pravesh Parekh, Diliana Pecheva, Anders M. Dale, Ole A. Andreassen, Paul M. Thompson, Christopher R.K. Ching, ENIGMA Bipolar Disorder Working Group

## Abstract

Compared to traditional gross volumetrics, surface-based models provide greater spatial precision for understanding brain alterations related to developmental, neurological, and psychiatric disorders. Large-scale brain initiatives are combining data from around the world to discover and improve illness-related brain markers. Here, we present a toolkit for 3D brain geometry analysis aimed at addressing key challenges facing large-scale neuroimaging studies. Our framework incorporates scalable methods for multisite data integration, site-specific confound correction, accelerated statistical modeling, interpretable machine learning, and interactive results visualization. The toolkit was tested on data from 21 independently collected study samples participating in the ENIGMA Bipolar Disorder Working Group (N = 3,373). Compared to traditional volume features, we show how subcortical shape measures can be combined across study sites to capture spatially complex differences between diagnostic groups and associations with common treatments. Statistical modeling was accelerated using the Fast and Efficient Mixed-Effects Algorithm (FEMA) and achieved a 16-fold reduction in computation time compared to traditional approaches. Machine learning models showed shape features may provide greater predictive performance over traditional volumes for both diagnostic and treatment prediction tasks, with interpretable weight maps providing insights into the local features driving model performance.

## I. Introduction

Altered subcortical volumes are commonly reported across studies of neurological [1], [2] and psychiatric illness [3], [4], [5]. Gross volumes are widely used, but more advanced surface-based and three-dimensional (3D) shape features offer spatial resolution sensitive to localized morphometric alterations not captured by gross estimates. Unlike neurodegenerative diseases, such as Alzheimer’s disease, where patterns of brain atrophy have been strongly linked to established underlying pathophysiology [6], psychiatric disorders such as bipolar disorder (BD) are associated with more subtle brain alterations, including smaller hippocampal and thalamic volumes that lack a clear link to underlying pathological processes [7]. Clinical factors such as symptom severity and treatment appear to influence the degree of brain variations in BD. Those with BD taking lithium treatment show patterns of larger brain volumes, whereas antiepileptic and antipsychotic medications have been associated with smaller volumes [3]. Such clinical factors can influence machine learning–based diagnostic performance trained on neuroimaging features in BD [8], and other psychiatric disorders [9]. A better understanding of the distribution of BD-related alterations across subcortical brain structures could drive insights into the vulnerabilities of particular subnuclei and how they result in disrupted brain networks driving BD symptoms [10]. Consequently, fine-grained mapping of treatment-related brain changes in large, multisite datasets may offer more clinically-relevant insights for predicting therapeutic response over time.

The Enhancing Neuro Imaging Genetics through Meta-Analysis (ENIGMA) Consortium [11] applies standardized data processing and analysis pipelines to independently collected neuroimaging samples from around the world to identify more replicable and generalizable brain markers in over 30 brain conditions. The ENIGMA Subcortical Shape Pipeline [12] is a standard framework for extracting vertex-wise radial distance and surface Jacobian measures across thousands of vertices spanning 7 subcortical structures including the hippocampus, amygdala, accumbens, caudate, putamen, pallidum, and thalamus. This pipeline has been applied to the largest international samples of schizophrenia [13], major depressive disorder [14], substance use disorders [15], Parkinson’s disease [2], and 22q11.2 copy number variation syndromes [16].

Building on this established framework, we introduce a Python-based 3D Brain Geometry Analysis Toolkit (Fig. 1) designed to facilitate large-scale, multi-site analyses of brain geometry data. The toolkit supports key steps to (1) integrate multisite samples, (2) perform efficient vertex-wise statistical modeling, (3) model potential site-related confounds, (4) facilitate interpretable machine learning analyses, and (5) generate static/interactive results visualizations. In this work, we evaluate the toolkit using subcortical shape data from the ENIGMA Bipolar Disorder Working Group (ENIGMA-BD) (N = 3,373). We assessed computational efficiency, the effects of data imputation methods, and the interpretability of machine learning models applied to surface-based features for both diagnostic and more clinical tasks.

**Fig. 1.**
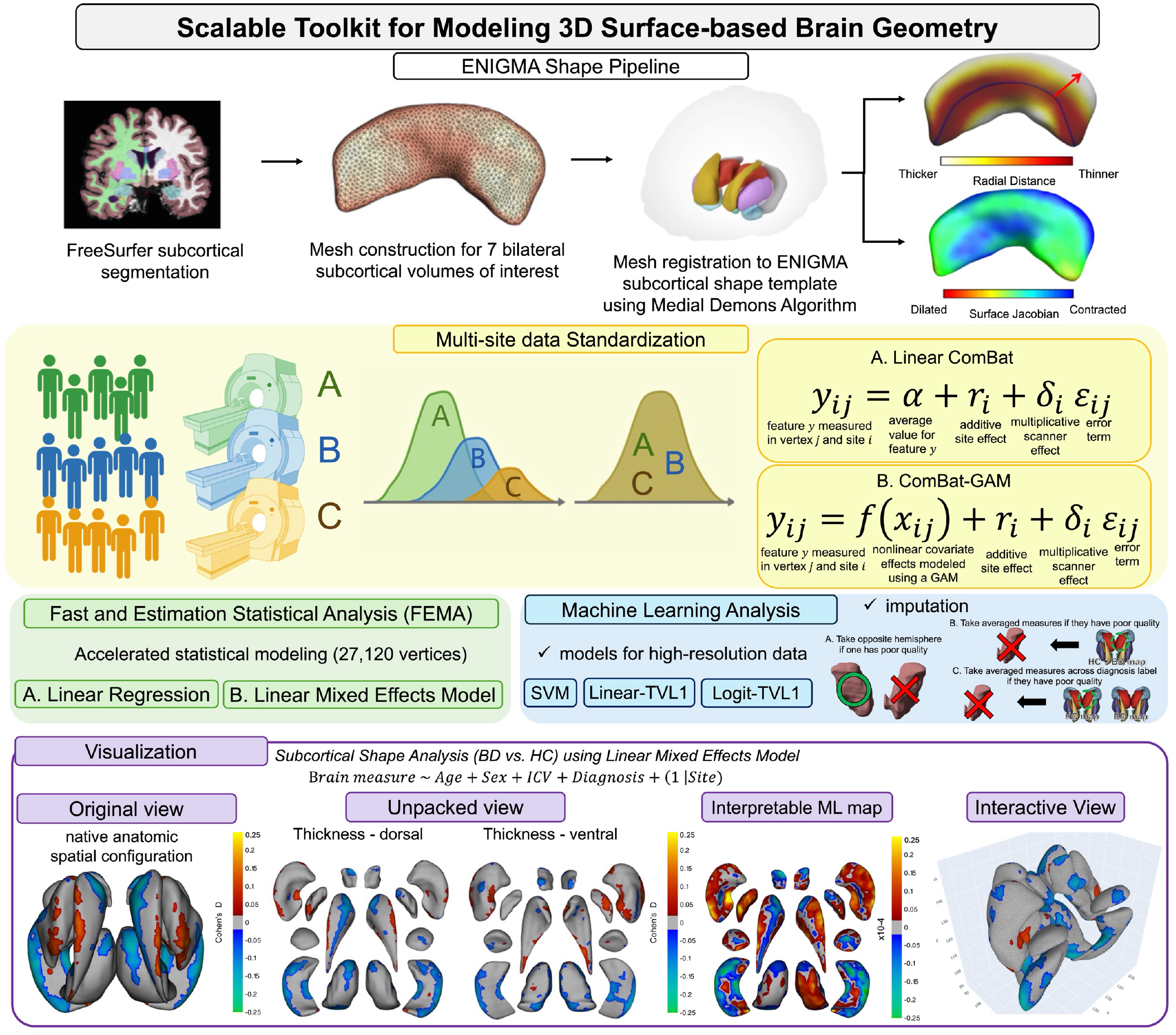
Python-based 3D brain geometry analysis toolkit overview

## II. Methods

### A. Data and Processing

3D T1-weighted brain MRI data from 21 independently collected ENIGMA-BD samples were included (BD=1,487, healthy controls (HC)=1,886; age: 8.0–83.0 years, 42% male). We first derived bilateral subcortical (gross) volumes (hippocampus, amygdala, caudate, putamen, pallidum, thalamus, and nucleus accumbens) using the ENIGMA-standard FreeSurfer (v5.3) protocol [17]. Next, the ENIGMA Subcortical Shape Pipeline [18] was used to extract vertex-wise shape features (radial distance or thickness) totaling 27,120 vertices from the 14 subcortical structures. All brain features underwent ENIGMA-standardized visual quality inspection by an expert rater to exclude poorly segmented shape models [19]. This same criterion was used in several published ENIGMA subcortical shape papers [13], [14], [20]. The 4% of cases failing quality control were excluded from subsequent statistical analysis.

**TABLE 1.** I.Demographic characteristics of ENIGMA-BD 21 sites’ samples.

| Site | # of BD/HC (% Male) | Age Mean (SD) |
| --- | --- | --- |
| Barcelona | 102/117 (46%) | 41.5 (±9.4) |
| BiDirect | 42/40 (43%) | 51.0 (±7.5) |
| Cardiff | 91/61 (33%) | 40.0 (±8.7) |
| CHRM2 | 42/54 (49%) | 41.5 (±13.0) |
| COGSBD | 59/31 (50%) | 37.1 (±11.2) |
| Halifax | 62/52 (38%) | 38.0 (±14.7) |
| Houston | 258/171 (43%) | 26.5 (±14.7) |
| IDIBAPS | 25/57 (43%) | 37.5 (±13.6) |
| Japan | 155/557 (36%) | 46.0 (±15.9) |
| KCL | 23/22 (31%) | 42.1 (±14.4) |
| Montpellier | 90/64 (40%) | 38.0 (±11.1) |
| MUSC | 68/26 (49%) | 38.7 (±9.3) |
| NUIG | 70/95 (56%) | 36.7 (±12.2) |
| Paris | 36/55 (51%) | 34.5 (±13.3) |
| Penn | 58/88 (45%) | 38.2 (±9.8) |
| SBA | 30/43 (44%) | 34.0 (±10.5) |
| Singapore | 43/44 (45%) | 23.4 (±3.9) |
| Sydney | 58/99 (41%) | 35.0 (±11.1) |
| Tulsa | 68/90 (29%) | 22.8 (±4.5) |
| UBC | 64/42 (47%) | 50.4 (±13.2) |
| VA San Diego | 43/78 (39%) | 37.8 (±11.6) |
| **Total** | **1487/1886 (42%)** | **38.0 (±15.0)** |

### B. ENIGMA Subcortical Shape Features

The ENIGMA Subcortical Shape Pipeline [18] represents each subcortical structure as a triangular surface mesh, with vertex coordinates defining three-dimensional geometry. Individual meshes are nonlinearly registered to a common ENIGMA template using the Medial Demons framework to obtain vertex correspondence across individuals [1]. Anatomical alignment is achieved by incorporating medial shape representations together with intrinsic geometric features to guide the registration process. This pipeline computes the following two measures:

#### Radial Distance (RD)

RD is a local thickness feature defined as the shortest Euclidean distance between a given surface mesh vertex and the medial curve c(t) of the structure. For a given vertex p∈M, this distance is computed as follows:

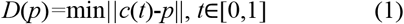

#### Tensor-Based Morphometry (TBM)

Local surface deformations are assessed using TBM, which quantifies the deformation needed to register an individual’s surface to the template [21]. This is achieved by computing a differential mapping between the tangent spaces of the template *M*_*t*_ and subject surfaces M, replacing the Jacobian:

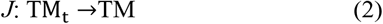

Here, the Jacobian matrix *J* describes the local linear transformation that maps a neighborhood on the template surface to the corresponding region on the subject’s surface, thereby capturing local geometric distortions in a surface-intrinsic manner. Full tensor-based analyses using the Jacobian determinant reflect the local surface expansion or contraction required to align the template with the subject, or the relative areal scaling between corresponding surface patches. For statistical analysis, the Jacobian determinant is log-transformed, yielding the log-Jacobian (log-Jac) feature, which is approximately normally distributed and provides a sensitive measure of localized morphometric variation. In this study, we focused primarily on the thickness (radial distance) shape measures for downstream statistical and machine learning analysis.

### C. Multisite Data Standardization

Our toolkit is equipped with ComBat and ComBat-GAM methods, which are widely used in multisite neuroimaging studies to adjust for potential site confounds using empirical Bayes, while preserving variability from biological factors including diagnosis, age, and sex. ComBat-GAM incorporates potential nonlinear age effects using generalized additive modeling. We adapted a Python-based source code for ComBat [22] and ComBat-GAM [23].

### D. Mass-Univariate Statistical Analysis

Our toolkit includes classic linear modeling features. Here, we tested 1) BD vs HC diagnostic differences and 2) associations between brain features and medication treatment at time of scan using both gross volumes and shape features. Here we report only results from the RD (local thickness) models, with Jacobian results to be included in a forthcoming analysis. All models were adjusted for age, sex, and intracranial volume (ICV). To control for potential site effects, we tested models including 1) raw brain measures with site included as a random effect, and 2) ComBat- and ComBat-GAM adjusted brain measures. All results were corrected for multiple comparisons (False Discovery Rate (FDR) *q*<0.05). Our toolkit also includes a module for efficient vertex-wise fixed- and mixed-effects modeling using the Fast and Efficient Mixed-Effects Algorithm (FEMA) [24]. FEMA fits linear mixed-effects models with fast runtime and low memory usage using sparse matrix algebra and parallel computation. Computational efficiency of FEMA and MATLAB *fitlme* were compared using downstream group effects analysis, while adjusting for the aforementioned covariates including site as a random effect. We calculated Cohen’s *d* (Fig. 2 and Fig. 3) using the *t* statistic from linear mixed-effects models, to interpret the relative effect sizes in line with our prior ENIGMA studies [25].

**Fig. 2.**
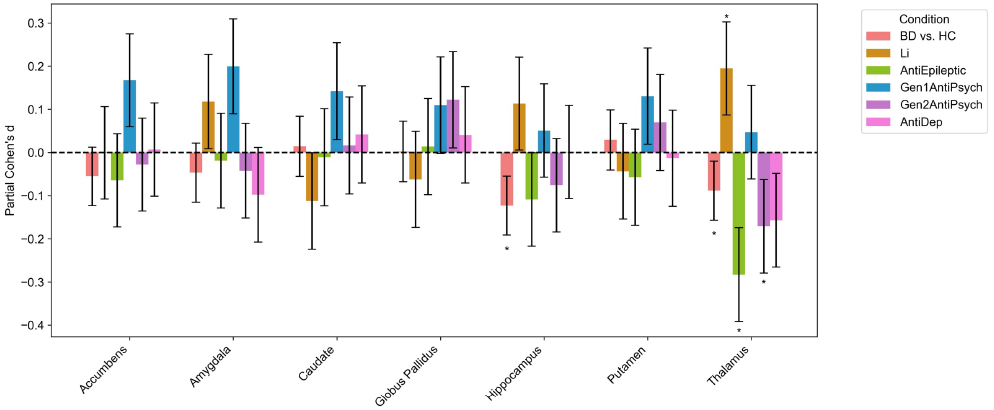
Cohen’s *d* effect sizes from gross volumetric analysis of BD versus HC group differences and association with medication time at scan. Asterisks indicate significant results after FDR correction (*q*<0.05).

**Fig. 3.**
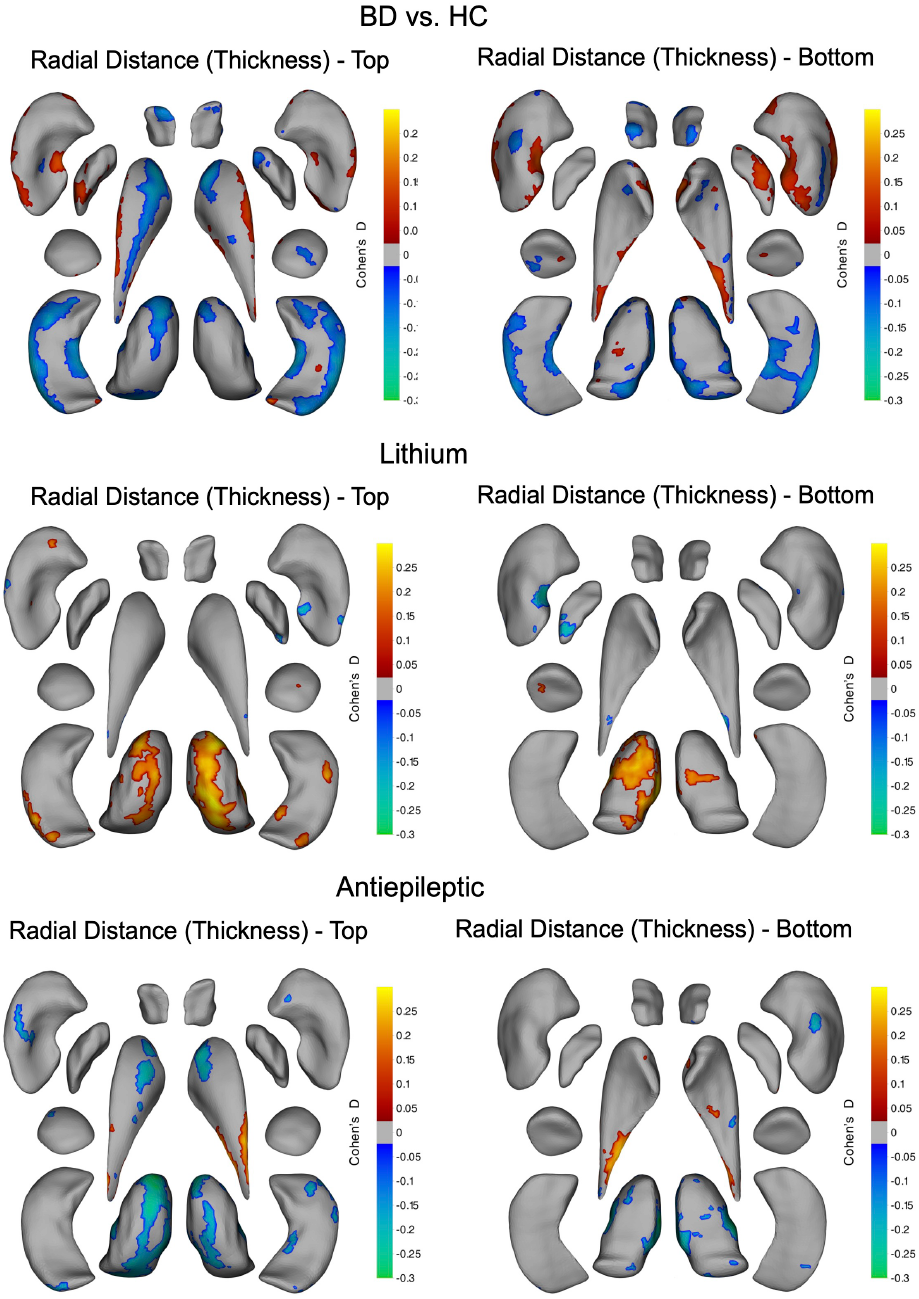
‘Unpacked’ results visualization showing Cohen’s *d* effect size maps from linear mixed-effects models adjusted for age, sex and intracranial volume (ICV) and site modeled as a random effect. Warmer colors indicate thicker regions (greater radial distance), and cooler colors indicate thinner regions associated with BD (compared to HC) and either lithium or antiepileptic treatment at time of scan. Gray regions indicate no significant difference or association after FDR correction for multiple comparisons.

### E. Machine Learning Functionality

The toolkit includes machine learning modules providing linear and logistic regression with L1 regularization for sparse solutions and a Total Variation (TV) term for spatial smoothness of vertex-wise coefficients. This dual-penalty regularization framework, referred to as Logit-TVL1 [26], learns discriminative weights that are associated with groups or clinical factors. These weights can be projected to the ENIGMA subcortical brain template to visualize stable and interpretable predictive brain regions. Logit-TVL1 has been widely applied in diffusion MRI studies of neurological disorders [27] and mesh-based regression and spectral analysis [28]. We applied this framework to BD versus HC diagnosis and medication prediction using the ComBat-GAM standardized features. A 10-fold cross-validated grid search was conducted to identify the optimal parameters, and model performance was evaluated using area under the receiver operating characteristic curve (AUC) and balanced accuracy (BAC). The significance of machine learning models’ performance was evaluated using a t-test on cross validation BAC.

### F. Imputation Methods

As missing data or exclusion after quality control procedures can influence machine learning model performance, our toolkit supports multiple imputation strategies, including:

(1)population-level imputation maps for each subcortical shape model based on the average shape metrics across all subjects,

(2)group-level imputation maps derived independently for BD and HC groups, (3) within individual hemisphere-flipping where poorly segmented structures in one hemisphere are replaced by properly segmented structures from that individual’s opposite hemisphere.

### G. Results Visualization

The toolkit includes a range of visualization functions for displaying statistical and machine learning results (Fig. 1), including representations that preserve neuroanatomical correspondence as well as unpacked dorsal and ventral surface projections. A 3D viewing tool allows users to interact with surface-based results via web browser (see https://yim-igc.github.io/ENIGMA_BD_Shape_LME_details/).

## III. Result

### A. Gross Volume and Subcortical Shape Alterations in BD

BD was associated with smaller gross hippocampal (Cohen’s *d*_BD-HC_ = −0.12) and thalamic volumes (*d*_BD-HC_ = −0.08) relative to HC (Fig. 2). Antiepileptic and antipsychotic treatment was associated with smaller thalamic volumes (*d*_AntiE_ = −0.28, *d*_Gen2AntiPsych_ = −0.17), whereas lithium treatment was associated with larger thalamic volumes when compared to individuals not taking those medications.

Shape analysis revealed complex and spatially distributed differences between BD and HC (Fig. 3). BD was associated with patterns of lower radial distance or thinner subregions of the hippocampus and thalamus. The putamen and caudate showed more complex patterns, with some areas displaying greater thickness alongside focally thinner regions in those with BD. Lithium treatment use was generally associated with patterns of greater thickness, especially in the thalamus when compared to those with BD not taking lithium. Antiepileptic use was mainly associated with the opposite pattern, or lower thickness in the thalamus and caudate. The putamen and caudate showed mixed patterns of both higher and lower thickness associated with the different treatment classes.

Compared to the traditional linear modeling approach (*fitlme*), FEMA resulted in a 16-fold reduction in computational time needed to fit the vertex-wise linear models.

### B. Machine Learning Analysis

Machine learning models trained on gross subcortical volume features did not show significant prediction performances for either BD diagnosis or medication at time of scan tasks (FDR *q*>0.05). However, TV-L1 models leveraging vertex-wise subcortical shape features did show significant balanced accuracy across all three tasks, including BD diagnosis: 55.4%, lithium treatment: 57.2%, and antiepileptic treatment: 55.2% (see models trained without imputation in Table 2).

**TABLE II.** The predictive performance OF Logit TV-L1 with different imputation strategy.

| Task | Imputation strategy | BAC | AUC |
| --- | --- | --- | --- |
| BD vs. HC | Without imputation | 55.4±1.9 | 55.4±1.9 |
|  | Single population-average imputation | 54.7±2.5 | 54.7±2.5 |
|  | Diagnosis-specific imputation | 54.6±2.6 | 54.6±2.6 |
|  | Hemisphere-flip imputation | 55.4±2.1 | 55.4±2.1 |
| Lithium prediction | Without imputation | 57.2±3.8 | 57.2±3.8 |
|  | Single population-average imputation | 55.2±4.7 | 55.2±4.7 |
|  | Diagnosis-specific imputation | 55.8±4.6 | 55.8±4.6 |
|  | Hemisphere-flip imputation | 55.1±5.2 | 55.1±5.2 |
| Antiepileptics prediction | Without imputation | 55.2±4.7 | 55.2±4.7 |
|  | Single population-average imputation | 54.6±5.1 | 54.6±5.1 |
|  | Diagnosis-specific imputation | 54.0±3.8 | 54.0±3.8 |
|  | Hemisphere-flip imputation | 54.2±5.5 | 54.2±5.5 |

Using a single population-average imputation map led to small reductions in performance across tasks (BD diagnosis BAC = 54.7; lithium prediction = 55.2; antiepileptic prediction = 54.6). Diagnosis-specific imputation produced a similar performance (BD diagnosis BAC = 54.6; lithium prediction = 55.7; antiepileptic prediction = 54.6). Hemisphere-flip imputation resulted in the same BD diagnosis performance (BAC = 55.4) as the non-imputed model and resulted in slightly worse lithium (BAC = 55.1) and antiepileptic (BAC = 54.2) prediction performance. AUC showed similar results to BAC (Table 2). Across all strategies, imputation led to slight reductions in predictive performance.

### C. Interpretable Machine Learning Maps

The spatial distribution of thickness differences observed in linear statistical analyses (Fig. 3) showed partial correspondence with regions driving machine learning performance (Fig. 4). The patterns of Logit-TVL1 weight maps depended on the regularization parameters determined by validation performance. In particular, increasing the TV penalty produced progressively sparser patterns, which corresponds to reduced spatial smoothing and results in weight maps that may be more challenging to interpret without arbitrary thresholding.

**Fig. 4.**
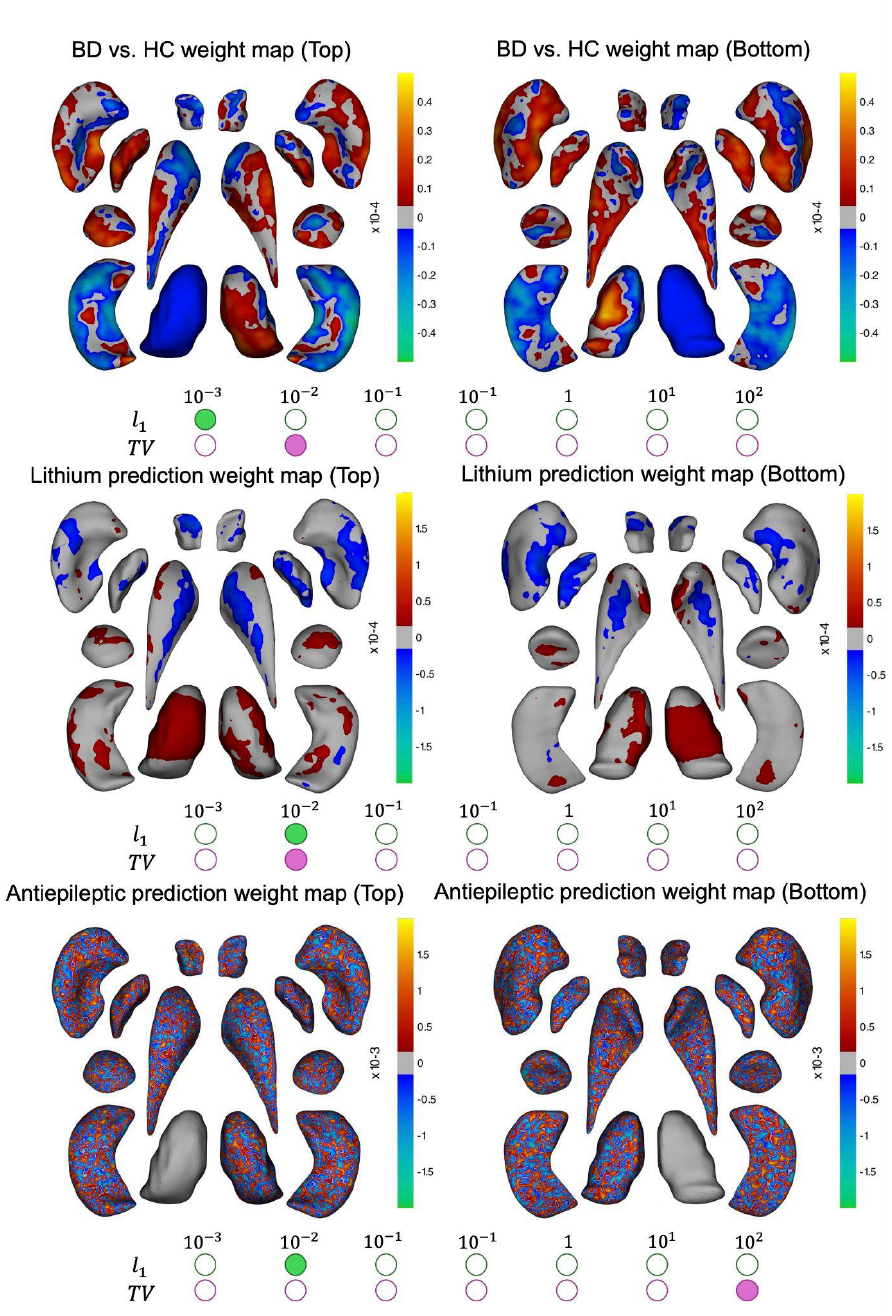
Weight maps from machine learning analysis using Logit-TVL1 trained with radial distance for three tasks: HC vs. BD classification, lithium and antiepileptic use at time of scan prediction.

## IV. Discussion & Conclusion

Here, we applied a scalable Python-based 3D brain geometry toolkit to the largest sample of subcortical volumes and shape features in those with BD to demonstrate multisite the feasibility of data integration, site correction modeling, missing data imputation procedures, accelerated mass-univariate statistical modeling, and interpretable machine learning to perform diagnostic and clinical prediction tasks.

Conventional vertex-wise mixed-effects analyses are often limited by excessive computational cost and memory demands, particularly in large consortium or biobank-scale datasets with thousands of individuals. In our benchmarking experiments, FEMA reduced statistical computation time by 16-fold relative to MATLAB’s *fitlme* [29], which will be essential for scaling toward high-dimensional neuroimaging analyses in large samples.

Whereas machine learning models based on traditional gross volume measures did not show significant performance in the selected tasks, the Logit-TVL1 models trained on subcortical shape features provided modest performance in classifying BD from HC as well as lithium or antiepileptic treatment at the time of scan. The Logit-TVL1 model predicted lithium or antiepileptic use at the time of scan slightly better than diagnosis of BD, which is consistent with prior studies based on volumetric measures [7], cortical thickness [3], and hippocampal subfield analyses [30].

Imputation analyses highlighted the challenges inherent to large-scale surface-based neuroimaging studies. Based on the current sample and modeling tasks, all imputation strategies resulted in small decreases in predictive accuracy. These findings may reflect the substantial inter-individual variability in subcortical morphology in BD, likely influenced by clinical factors such as illness duration, severity, treatment and other environmental factors.

Our framework includes some limitations currently under development. The present toolkit implementation focuses on subcortical shape features. Work is underway to extend functionality to cortical surface representations, enabling unified analyses of cortical and subcortical geometry. Second, although Logit-TVL1 provides interpretable spatial patterns of task performance, it represents only one class of explainable models. Forthcoming toolkit features will include spherical convolutional neural networks [31] and mesh-based vision transformers [32], which may better capture higher-order geometric relationships. Finally, integrating surface-based morphometric features with complementary biological data, such as polygenic risk scores, gene expression profiles, cytoarchitectural maps, and neurotransmitter distributions, represents an important next step toward multiscale modeling of BD and related conditions.

In summary, our toolkit to address key challenges facing large-scale studies of 3D brain geometry and will help to standardize multisite data integration, imputation, efficient statistical modeling, interpretable machine learning, and interactive results visualizations to facilitate the discovery of more reliable and clinically-actionable brain signatures across disorders.

## Acknowledgment

This work was made possible by the following additional ENIGMA Bipolar Disorder Working Group Members: Ana M. Diaz Zuluaga, Andriana Karuk, Annabella Di Giorgio, Benson Mwangi, Boris Gutman, Bronwyn Overs, Carlos López Jaramillo, Colm McDonald, Dan Stein, Dara M Cannon, David Glahn, Diego Hidalgo-Mazzei, Dominik Grotegerd, Edith Pomarol-Clotet, Eduard Vieta, Enric Vilajosana Chertó, Fabio Sambataro, Fleur Howells, Freda Scheffler, Geraldo Busatto, Gerard Anmella, Giovana B. Zunta-Soares, Gloria Roberts, Henk Temmingh, Ian Gotlib, Ingrid Agartz, Jair C. Soares, James A. Karantonis, James Prisciandaro, Janice M. Fullerton, Joaquim Radua, Joaquim Radua, Jonathan Savitz, Josselin Houenou, Kang Sim, Kenichiro Harada, Klaus Berger, Koji Matsuo, Lakshmi Yatham, Lars Tjelta Westlye, Leila Nabulsi, Lisa Eyler, Lisa Furlong, Luisa Klahn, Marco Hermesdorf, Marcus V. Zanetti, Matthew D. Sacchet, Matthew Kempton, Mikael Landen, Mon-Ju Wu, Ole A. Andreassen, Emilie Olie, Pedro Rosa, Philip Mitchell, Raymond Salvador, Rayus Kuplicki, Salvador Sarró, Sophia I. Thomopoulos, Susan Rossell, Tamsyn Van Rheenen, Theodore Satterthwaite, Tilo Kircher, Tomas Hajek, Udo Dannlowski, Xavier Caseras. This work was supported by NIH grants R01MH129742, AG081571 and R21MH139001, and R56 AG058854. The BiDirect was supported by grants of the German Ministry of Research and Education (BMBF) to the University of Muenster (01ER0816 and 01ER1506). This work was funded by the German Research Foundation (DFG, grant FOR2107 DA1151/5-1, DA1151/5-2, DA1151/9-1, DA1151/10-1, DA1151/11-1 to UD; SFB/TRR 393, project grant no 521379614) and the Interdisciplinary Center for Clinical Research (IZKF) of the medical faculty of Münster (grant Dan3/022/22 to UD). The MUSC was funded by K23 AA020842. The Houston cohort was partially supported by NIMH (1R01MH085667), John S. Dunn Foundation (Houston, Texas), and Pat Rutherford Chair in Psychiatry (UTHealth Houston). The Paris cohort was funded by the French Agence Nationale Pour la Recherche. The Sydney (UNSW) study was supported by the Australian National Medical and Health Research Council (NHMRC) Program Grant 1037196 to Philip B. Mitchell, NHMRC Project Grants 1066177 and 1063960 to Janice M. Fullerton, NHMRC Investigator Grants 1176716 to Peter R. Schofield and 1177991 and Philip B. Mitchell, and NHMRC & Medical Research Futures Fund Grant 1200428 to Janice M. Fullerton. Janice M. Fullerton is the grateful recipient of the Janette Mary O’Neil Research Fellowship. Additional philanthropic support was provided by the Lansdowne Foundation, GoodTalk charity, the Gordon Pettigrew Family, Mrs Betty C. Lynch OAM (dec) and the Aberdeen Foundation. Pravesh Parekh was supported by the European Union’s Horizon 2020 research and innovation programme under the Marie Skłodowska-Curie grant 801133; Research Council of Norway grant 324252; NIH grants U24 DA041123 and U24 DA055330. Anders M. Dale is supported by NIH grants U24 DA041123, DA055330, R01 AG076838, and OT2 HL161847. Dr. Tamsyn Van Rheenen was supported by an Al and Val Rosenstrauss Fellowship from the Rebecca L Cooper Medical Research Foundation. James A. Karantonis was supported by Swinburne University/an Australian Postgraduate Award. Lisa Furlong was supported by the Australian Rotary Health and the Ian Parker Bipolar Research Fund; Brunslea Park Estate. Susan Rossell was supported by an NHMRC Senior Fellowship. The GIPSI cohort was supported by PRISMA UT and MINCIENCIAS. The COGSBD study was financially supported by the NHMRC (1060664), Henry Freeman Trust, Jack Brockhoff Foundation, University of Melbourne, Barbara Dicker Brain Sciences Foundation, Rebecca L Cooper Foundation and the Society of Mental Health Research. The authors acknowledge the facilities and scientific assistance of the National Imaging Facility, a National Collaborative Research Infrastructure Strategy (NCRIS) capability, at the Swinburne Neuroimaging Facility, Swinburne University of Technology. This work was supported by MH083968 from the US National Institutes of Health and the Desert-Pacific Mental Illness Research Education and Clinical Center.

